# Downregulation of *Yap1* During Limb Regeneration Results in Defective Bone Formation in Axolotl

**DOI:** 10.1101/2022.06.12.495807

**Authors:** Sadık Bay, Gürkan Öztürk, Nesrin Emekli, Turan Demircan

## Abstract

The Hippo pathway plays an imperative role in cellular processes such as differentiation, regeneration, cell migration, organ growth, apoptosis, and cell cycle. Transcription coregulator component of Hippo pathway, YAP1, promotes transcription of genes involved in cell proliferation, migration, differentiation, and suppressing apoptosis. However, its role in epimorphic regeneration has not been fully explored. The axolotl is a well-established model organism for developmental biology and regeneration studies. By exploiting its remarkable regenerative capacity, we investigated the role of *Yap1* in the early blastema stage of limb regeneration. Depleting *Yap1* using gene-specific morpholinos attenuated the competence of axolotl limb regeneration evident in bone formation defects. To explore the affected downstream pathways from *Yap1* down-regulation, the gene expression profile was examined by employing LC-MS/MS technology. Based on the generated data, we provided a new layer of evidence on the putative roles of increased protease inhibition and immune system activities and altered ECM composition in diminished bone formation capacity during axolotl limb regeneration upon *Yap1* deficiency. We believe that new insights into the roles of the Hippo pathway in complex structure regeneration were granted in this study.

## 1. INTRODUCTION

The Hippo pathway regulates multiple cellular processes, including organ growth, cell proliferation, apoptosis, the transmission of mechanical signals, cell cycle, cell migration, differentiation, organ and limb regeneration (Hayashi et al., 2015; Moya and Halder, 2018; Panciera et al., 2017). This pathway also cross-talks with other signaling pathways such as Wnt (Wingless/Ints), GPCRs (G-Protein Coupled Receptors), RTKs (Receptor Tyrosin Kinases) (Moya and Halder, 2018) to function in organ development and homeostasis. Accumulated studies in mammals and highly regenerative model organisms have demonstrated that misregulation of the Hippo pathway affects a wide range of biological processes and pathology of the diseases.

In addition to its regulatory roles during embryonic development, the Hippo signaling pathway controls organ size maintenance following the tissue or organ regeneration in animals. During mouse embryogenesis, deletion of the *MST1* and *MST2* kinase genes showed enlargement of the liver and heart (Camargo et al., 2007; Dong et al., 2007). Previous studies have suggested that the apparent diminish in cardiac regeneration capacity of mammals a few days after birth is due to the decrease in the transcriptional activity of *YAP1*, which is negatively regulated by increased activity of the tumor suppressor proteins LATS1 and LATS2 (Heallen et al., 2013; Xin et al., 2013). In another study conducted in adult mice, a high level of *YAP1* was associated with liver enlargement (Benhamouche et al., 2010). In mammals, albeit with limited regeneration capacity, there is accumulating evidence for the induction of tissue repair upon *YAP/TAZ* activation in different tissues. Previous findings have linked intestinal regeneration with *YAP1* activity. After an intestinal injury, the inhibitory phosphorylation of *YAP1* by the upstream kinases is lifted, and the active *YAP1* is translocated to the nucleus. After this localization, the transcriptional activity provided by *YAP1* initiates the stem cell self-renewal programming directed by the Wnt pathway (Gregorieff et al., 2015; Yui et al., 2018). Moreover, enrichment of *YAP1* in the LGR5+ stem cell compartment of the intestinal crypt during homeostasis and its upregulation during regeneration (Cai et al., 2010) highlighted the active participation of *YAP1* in these processes. As in the intestine, the regeneration program in the liver requires activation of *YAP1*, which is evident by the enhancement of liver regeneration consequent to experimental activation of *YAP1* (Bai et al., 2012), even in aged mice with an impaired liver regeneration potential (Loforese et al., 2017). Increased proliferation of embryonic cardiomyocytes following *YAP1* activation resulted in a continuous proliferation of cardiomyocytes in adults, which might be the key to enabling heart function restoration after infarction (von Gise et al., 2012). In addition to these organs, *YAP1* also plays essential roles in skin regeneration, as evident in the previous studies where the silencing of *YAP/TAZ* gene expression interferes with skin cell proliferation and successful skin regeneration in mammals (Johnson and Halder, 2014; Juan and Hong, 2016). The observed regulatory functions of *YAP1* in flatworms’ whole-body regeneration highlights the necessity of the *YAP1* activity in the regeneration process of invertebrates (Demircan and Berezikov, 2013; Lin and Pearson, 2014). Furthermore, studies in different model systems have exhibited that the Hippo pathway role in organ regeneration is highly conserved among animals (Hayashi et al., 2015).

Amphibians, unlike other tetrapods, display a higher organ and tissue regeneration ability (Nye et al., 2003) extended to complete regeneration of many organs and extremities such as the cornea, lens, retina, heart, spinal cord, brain, tail, and limb (Stocum, 2006). Axolotl, a fruitful model organism for developmental biology and regeneration studies, has a high regeneration capacity throughout its life due to larval-like characteristics in adulthood as it can not undergo metamorphosis naturally (Galliot and Ghila, 2010). Therefore, it is a widely utilized model to study the molecular mechanisms of regeneration, particularly to explore the key steps of and regulators in limb regeneration. Furthermore, as another remarkable experimental advantage, axolotls can be induced to metamorphosis by thyroid hormone administration, which allows researchers to evaluate the limb regeneration capacity at different developmental stages during adulthood (Rosenkilde et al., 1982). Initial reports indicating the drastic drop of regeneration capacity and fidelity upon metamorphosis suggest using pre-and post-metamorphic axolotls to expand our knowledge on limb regeneration (Demircan et al., 2018; Monaghan et al., 2014).

One of the critical cellular processes in organ regeneration is the regeneration’s size control step. In axolotl and various frog species, organs return to their original size after damage (Beck et al., 2009; McCusker et al., 2015; Slack et al., 2008). As documented before, differentiated cells turn into progenitor and stem cells and accumulate at the injury site upon an injury or amputation. The formation of regeneration-specific tissue, blastema, is indispensable to renew the missing parts. Later stages of regeneration can be summarized as restoration of the limb structure and reshaping of the newly formed limb. It has been described that the Yap protein is highly active during tail and limb regeneration in *Xenopus laevis*. YAP and TEAD4 proteins are essential to maintain original size, and the absence of YAP or TEAD4 proteins during regeneration leads to a shorter tail in tadpoles (Hayashi et al., 2014a). In addition to tail regeneration, Yap protein activation was detected during *Xenopus laevis* limb regeneration (Hayashi et al., 2014b). The signaling cascades regulating renewed structure’s final size in axolotl limb regeneration are not fully understood yet.

In this study, we investigated the putative roles of *Yap1* in axolotl limb regeneration. Significantly higher *Yap1* expression at mRNA and protein levels in neotenic axolotls compared to the metamorphic animals at 10 day-post amputation (dpa) prompted us to down-regulate the neotenic *Yap1* expression at the early blastema stage of limb regeneration. Microscopic, macroscopic and computational tomography (CT) based analyses revealed the defective bone regeneration due to the *Yap1* inhibition. To provide molecular clues, the proteome of *Yap1* inhibited and control animals were compared. We identified essential peptidase activity, immune system, and wound healing related pathways enriched by the differentially expressed proteins between two groups. The findings of this study will contribute to the understanding of the *Yap1* roles in successful regeneration.

## 2. Material and Methods

### 2.1 Animal Husbandry and Ethical İssues

Axolotls 12-15 cm in size and 1 year old were used in this study. Animals were maintained in Holtfreter’s solution at 18±2 °C temperature as one individual in an aquarium as described in a previous study (Demircan et al., 2019) throughout the experiments. Required permission and approval for this study was obtained from the local ethics committee of animal experiments of Istanbul Medipol University (IMU-HADYEK, Approval Number: 38828770-604.01.01-E.14550).

### 2.2 Induction of Metamorphosis

A previously established protocol (Demircan et al., 2018) was followed to induce the metamorphosis for neotenic axolotls. Briefly, randomly selected animals (n=18) were treated with T3 (3,3’,5-Triiodo-L-thyronine – Sigma T2877) twice a week in Holtfreter’s solution with a 25 nM final concentration. Morphological alterations such as weight loss, disappearance of the fin and gills were monitored carefully to follow the transition to the metamorphic stage, and approximately after 4 weeks of treatment, the T3 concentration was reduced to half. The hormone was administered for an additional two weeks. After fully metamorphosis was accomplished, animals were kept for a month without hormone treatment to allow them to adapt to the terrestrial life conditions.

### 2.3 Sample Collection

Prior to amputations, the animals were anesthetized using 0.1% Ethyl 3-aminobenzoate methanesulfonate (MS-222, Sigma-Aldrich, St Louis, MO, USA) dissolved in Holtfreter’s solution. Right forelimbs of animals were amputated at mid zeugopod level under a dissecting microscope (Zeiss - StereoV8). To compare *Yap1* mRNA levels between neotenic (n=15) and metamorphic animals (n=15), tissue samples were collected at the early steps of regeneration (1, 7,10,14,21 dpa). Blastema samples of neotenic (n=3) and metamorphic (n=3) axolotls at 10 dpa were obtained to evaluate the YAP1 protein expression level by immunohistochemistry. The efficiency of morpholino injections was assessed on blastema samples (n=3 for *Yap1* and n=3 for control morpholino injections) removed at 16 dpa. For proteomics and qRT-PCR experiments blastema samples were collected at 16 dpa for *Yap1* and control morpholino injected animals (n=12 for each group). All collected tissue samples according to the defined time points were cryopreserved in liquid nitrogen and stored at −80°C until RNA or protein isolation.

### 2.4 *Yap1* Morpholino Design and Administration

GeneTools (https://www.gene-tools.com/) was used to design the morpholino oligo (MO) (5’-CCTCTTACCTCAGTTACAATTTATA-3’) for *Yap1* inhibition. As a negative control, a standard oligo sequence (5’-CCTCTTACCTCAGTTACAATTTATA-3’) suggested by GeneTools was used. Morpholinos were dissolved in 2X PBS with a final concentration of 500 μM. To assess the efficiency of morpholinos, each morpholino was injected into 3 axolotls at 10 dpa. To determine the effect of morpholinos on regeneration, each morpholino was administrated to another 9 axolotls at 10 dpa. For proteomics and qRT-PCR experiments each morpholino was injected into 12 axolotls at 10 dpa. Injection into blastema using Hamilton injector (SGE 025RN, 25 GA 50MM) was followed by electroporation as described elsewhere (Leigh et al., 2020). Electroporation efficiency was assessed by co-injection of GFP encoding plasmid. Fluorescent imaging of blastema tissues was performed using AxioZoom V16 (Zeiss) microscope before and on the 4th and 7th day after injection.

### 2.5 Quantitative Real-Time Polimerase Chain Reaction (qRT-PCR)

To compare the expression level of *Yap1* in neotenic and metamorphic axolotl blastema, tissue samples at 1, 7, 10, 14, and 21 were isolated (n=15 for neotenic and metamorphic animals, and n=3 per time point). To evaluate the efficiency of *Yap1* knockdown, blastema tissues were obtained 6 days after injection (n=3 for *Yap1* and control morpholinos). The details of the following qRT-PCR protocol was described elsewhere (Sibai et al., 2019). Briefly, RNA from collected samples was isolated using the Total RNA Purification Kit (Norgen-Biotek, 37500) according to the manufacturer’s protocol. The quality of isolated RNA was checked on 1% agarose gel. The quantity of RNA was measured using the nanodrop. ProtoScript First Strand cDNA Synthesis Kit (NEB, E6300S) was employed following the producer’s procedure to synthesize the complementary DNA starting with 1 μg total RNA. SensiFAST™ SYBR® No-ROX Kit (Bioline, BIO98005) was utilized in qPCR, considering the manufacturer’s suggestions. Primer sequences to amplify *Yap1* and *Elf1α* (housekeeping gene used for normalization) were provided in Table S1. qPCR experiments were performed in 3 biological and 3 technical replicates and CFX Connect-Real Time System (BIO RAD) device was used for this reaction. With 2^−ΔΔ*Ct*^ method, relative messenger RNA (mRNA) expressions were calculated.

### 2.6 Immunofluorescent (IF) Staining

To label the YAP protein in axolotl blastema tissues, a protocol used in *Xenopus laevis* (Hayashi et al., 2014b) was slightly modified and used. Blastema tissues (n=3 for neotenic and metamorphic animals) were first embedded in a tissue freezing medium (Leica, 14020108926). Then, the embedded tissues were placed on a Cryostat device (Leica, CM1950) for sectioning to get the slices with 25 μm thickness. The sections were placed on positively charged slides (Superfrost Plus, ThermoScientific, J1800AMNZ) and outlined with a PAP-PEN (Invitrogen, 008877). Later, sections were fixed 2 times with 4% PFA (Sigma, 158127) for 15 minutes followed by subsequent incubation steps in 0.1% Triton TX-100 (Sigma, T8787) for 20 minutes, 0.3% H_2_O_2_ (Sigma, H1009) for 10 minutes, and 2% FBS (ThermoFisher Scientific, 16140071) for 1 hour. Primary Rabbit-anti-YAP antibody (Cell Signalling Technology - 4912) diluted 1:250 was added to cover the samples on slides and an overnight incubation at +4 ^o^C was followed. Samples were incubated with Alexa Fluor 594 attached Goat-anti-Rabbit secondary antibody (Invitrogen, A11037, diluted 1:250) for 3 hours at room temperature. After the DAPI (Sigma, D9542) with 1:1000 dilution was added to stain the nucleus, slides were covered with the mounting solution (Mounting Medium, C9368). For imaging, a confocal microscope (Zeiss LSM800 with AiryScan) was used. As a negative control, slides stained with secondary antibody only were imaged. Antibody positive cells in tissues were counted with ZenBlue (Zeiss) software.

### 2.7 Computational Tomography (CT)

U-CT (MiLabs) device was used to determine whether bone development was affected by *Yap1* inhibition during limb regeneration (n=9 for control and *Yap1* morpholino injections). After the axolotls were anesthetized, they were placed in a tomography container suitable for the size of the analyzed animals. High-resolution scan mode was selected for imaging, and the ‘Reconstitution (MiLabs)’ software of the device was utilized to convert the images to 3D. The data was transferred to the ‘IMALYTICS Preclinical 2.1 (MiLabs)’ software for further processing to adjust the brightness and contrast.

### 2.8 Proteomics Analysis by LC-MS

Animals (n=24) were randomly divided into two groups and amputated as described above. At the early blastema stage (10 dpa), *Yap1* and control morpholinos were administrated to Yap1_KD or the control groups following the procedure described before. One week after the morpholino injections, 9 blastema tissues for each treatment were collected and gathered to form three replicates, in which three samples were pooled. For proteomics analysis, previously established protocols were followed (Cevik et al., 2016; Sibai et al., 2020). Briefly, mortar and pestle were used to powder the frozen blastema samples for protein extraction. Proteins were isolated using an ultrasonic homogenizer with three 5s on-5 s off cycles and quantified by Qubit 4.0. Filter-aided sample preparation (FASP) (Abcam - ab270519) method was applied to obtain the tryptic peptides, and the mixture of 500 ng tryptic peptides was used in triplicates in downstream steps. ACQUITY UPLC (Waters) chromatography coupled high-resolution mass spectrometry SYNAPTG2-Si (Waters) system was used for the analysis of peptides. The parameters defined in previous studies (Cevik et al., 2016; Sibai et al., 2020) were followed for peptide separation and MSE data collection. Obtained signals were preprocessed with ProteinLynx Global Server (v2.5, Waters) and a previously generated axolotl database (Demircan et al., 2017) was used to identify and review the peptide sequences. Progenesis QI for proteomics (v.4.0, Waters) software was employed for protein identification and quantitative analysis of detected peptides. Normalization of data was conducted according to the relative quantitation of non-conflicting peptides. ANOVA test was applied to filter the proteins with a p-value ≤ 0.01 and a 2-fold or greater differential expression level between the two conditions. To validate the proteomics results and evaluate the expression level of the genes with osteogenic or chondrogenic activity, 6 blastema tissues the RT-qPCR method described above was applied using the gene-specific primers (Table S1) Yap1_KD treatment (n=3) and control samples (n=3).

### 2.9 Enrichment of Gene Ontology (GO) Terms and KEGG Pathways

Differentially enriched (DE) proteins between the *Yap1* morpholino treated and control groups were tested for identifying Molecular function (MF), biological process (BP), and cellular component (CC) GO terms using “clusterProfiler’’ package in R (Yu et al., 2012a). The same package was used to explore the Kyoto Encyclopedia of Genes and Genomes (KEGG) pathways enriched by DE proteins. The parameters applied for enrichment analyses were as follows: p-value and q-value cutoffs were 0.05, and adjusted p-value was Benjamini & Hochberg (BH). Enriched GO terms and pathways were visualized on dot plot and bar plot graphs using the “ggplot2” package in R.

### 2.10 Statistics

GraphPad Prism 8 (San Diego, CA, USA) was used for statistical analysis. In order to test the normality of data distribution, the Shapiro-Wilk test was carried out. One-way ANOVA with post-hoc Tukey’s test was applied on qRT-PCR and proteomics data. The chi-squared test was employed in the percentages calculated of IF results. The student’s t-test was used to compare the limb measurement results. All p-values smaller than 0.05 were considered significant. An asterisk (*), two asterisks (**) or three asterisks (***) were used to indicate the significance of p values as follows: 0.05≤p-value≤0.01, 0.01<p-value≤0.001 or p<0.001, respectively.

## 3. RESULTS

### 3.1 Neotenic *Yap1* mRNA and Protein Levels Were Higher Than Metamorphic *Yap1* in Blastema

Considering the decreased regeneration capacity in metamorphic axolotls, we first compared the *Yap1* level in neotenic and metamorphic blastema tissues at 1, 7, 10, 14, and 21 dpa to inspect the putative link between *Yap1* and limb regeneration potential by employing the qRT-PCR (Fig. S1, Fig. 1A). Although a higher expression level of *Yap1* was detected in neotenic tissues compared to metamorphic ones at all time points, the greatest expression level difference was identified at 10 dpa. *Yap1* was substantially and significantly upregulated in neotenic blastema tissue compared to metamorphic one at mRNA level (>70x) (Fig. 1A). This finding was further validated by IF results, where the rate of *Yap* positive cells in neotenic blastema tissue was higher than the metamorphic samples (Fig. 1B) (13,12% and 3.03%, respectively). According to these results, *Yap1* levels were elevated at both mRNA and protein levels in neotenic animals in comparison to metamorphic ones at the early blastema stage of axolotl limb regeneration.

**Fig. 1.**
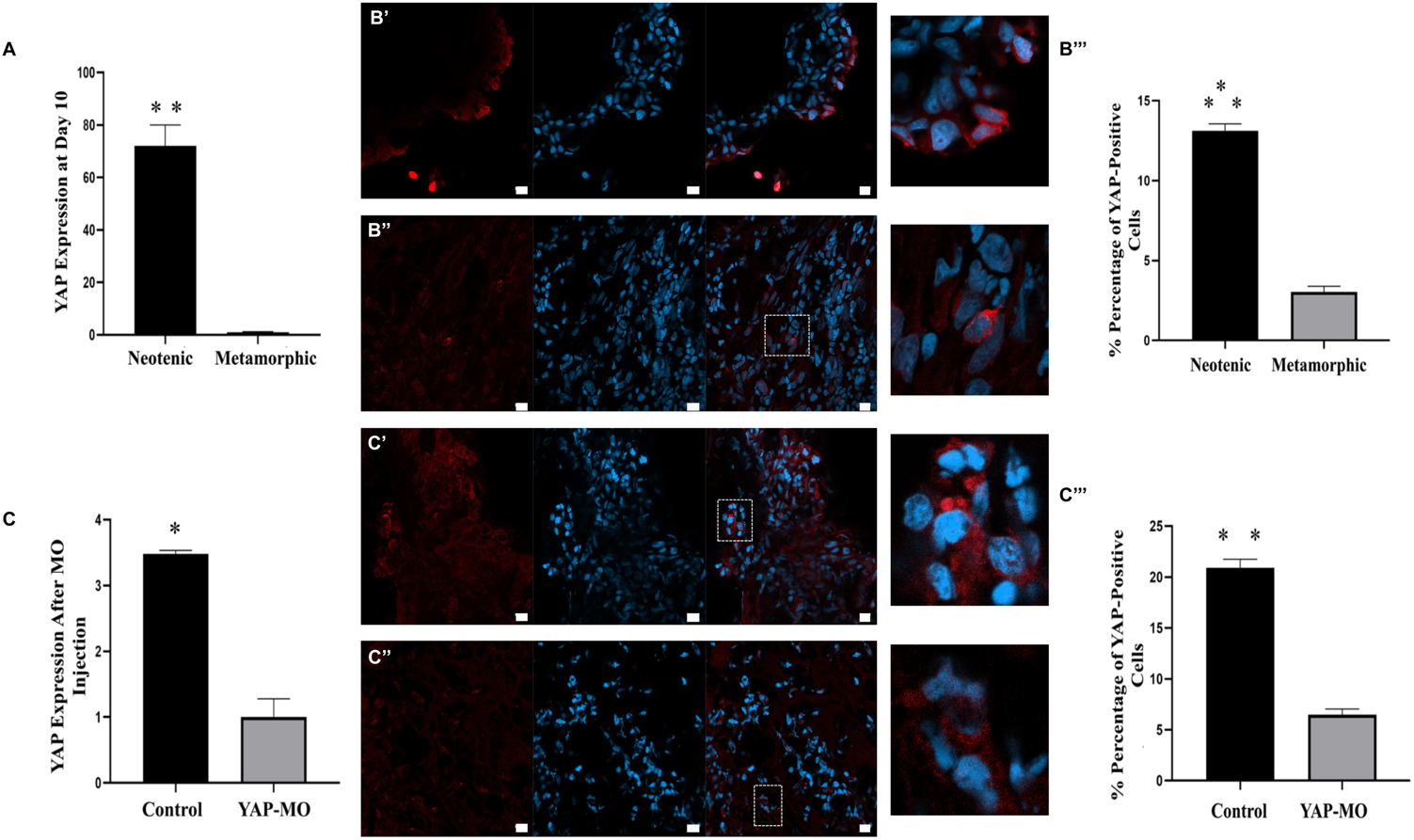
*Yap1* was downregulated during axolotl limb regeneration using morpholino (MO). Neotenic axolotl *Yap1* mRNA expression levels were 72 times more compared to metamorphic *Yap1* mRNA expression at 10 days post amputation (dpa) (A) (0.01<p**≤0.001). For determining YAP1 protein level, neotenic (B’) and metamorphic (B’’) blastema tissues were stained with YAP1 antibody at 10 dpa (B). Percentage of YAP1-positive cells was 13,12% in neotenic axolotl while percentage of YAP1-positive cells was 3,03% in metamorphic axolotl at 10 dpa (B’’’) (p***<0.001). YAP-MO were synthesized for inhibition of axolotl *Yap1* mRNAs. After MO injection to blastema tissue of neotenic axolotl at 10 dpa, qRT-PCR analysis were performed at 16 dpa. *Yap1* mRNA expression in YAP-MO animals decreased 3,48 times compared to control animals (C) (0.05≤p*≤0.01). Yap1 in control axolotl (D’) and Yap-MO (D’’) axolotl were imaged in blastema tissues (D). Yap-MO caused the level of Yap1 protein to decrease from 20,89% to 6,49% at 16 dpa (D’’’). Scale bars represent 20 μm.

### 3.2 An Efficient Knock-down of *Yap1* was Achieved by Morpholino İnjection

To inhibit the *Yap1* activity in blastema tissue, its mRNA was depleted using a designed gene-specific morpholino. The efficiency of morpholino on *Yap1* expression was tested by qRT-PCR and IF methods (Fig. 1C and 1D). Morpholinos were injected at 10dpa and mRNA level, and the rate of YAP+ cells was evaluated 2 and 6 days after administration of morpholino, respectively. The Yap-MO injected group (Yap1_KD) had a significantly reduced *Yap1* mRNA level (3.48x) compared to the control-MO injected group (Fig. 1C). The rate of YAP-positive cells in the Yap-MO injected group was detected significantly lower than the control-MO injected group (Fig. 1D) (6.49% and 20.89%, respectively). Hence, we concluded that Yap-MO was effective in inhibiting the *Yap1* activity, and we, therefore, used this MO in subsequent experiments.

### 3.3 *Yap1* Inhibition Interfered with Successful Bone Regeneration in Axolotl

After the Yap-MO injection animals were followed for long-term macroscopic and microscopic observations. Blastema measurements at 1-week post-injection indicated a significant decrease in blastema size between Yap-MO and control-MO injected animals (Fig. 2A and Fig. 2B). Macroscopic observations did not unveil any detectable morphological differences between the groups at 52 dpa (Fig. 2A). However, observed softer digit formation in the renewed limb prompted us to analyze the bone structure employing the Micro-CT at 52 dpa. Bone defects due to bone orientation disorder and bone loss were detected for YAP-MO injected axolotls (Fig. 2C), highlighting the roles of YAP1 protein in successful bone development and regeneration in the axolotl.

**Fig. 2.**
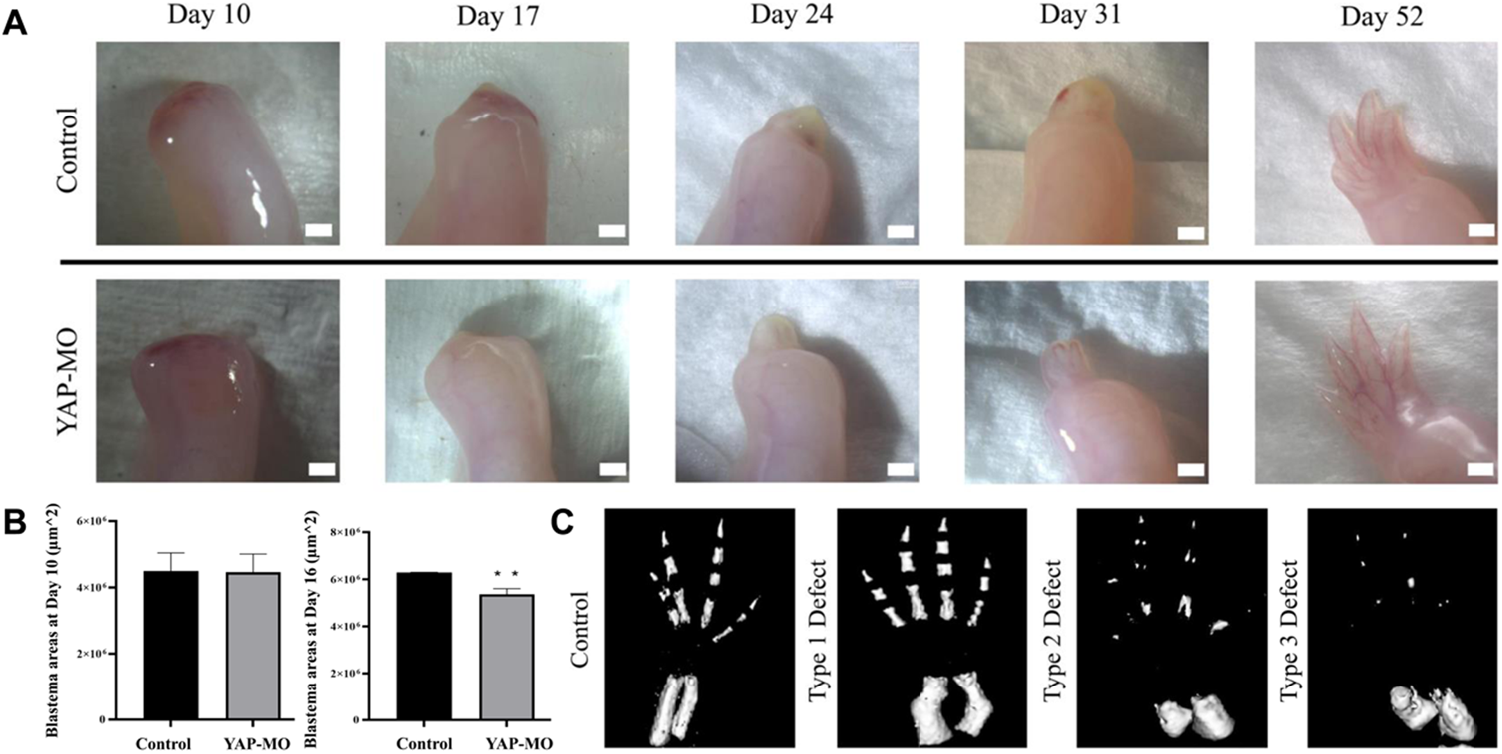
Yap-MO caused bone defects during axolotl limb regeneration. After Yap-MO injection, macroscopic imaging’s were performed at 10, 17, 24, 31 and 52 dpa. Different regeneration patterns were observed between control axolotls (n=3) and Yap-MO axolotls (n=3) as a result of long-term bright field imagings (A). Scale bars represent 1 mm. Yap-MO injection induced the decrease of blastema areas at 16 dpa. At 10 dpa, significant change was not determined between control axolotls (n=3) and Yap-MO axolotls (n=3). Blastema areas were decreased in Yap-MO axolotls (n=3) compared to control axolotls (n=3) (0.01<p**≤0.001) (B). Macroscopic differences were not detected in the bright-field results after regeneration was completed. Micro-CT analysis was performed to determine whether there was a defect in bone development after Yap-MO injection at 52 dpa. Bone orientation defect and bone loss defect were imaged with Micro-CT (C).

### 3.4 Proteomics Results Highlighted That *Yap1* Down-regulation Altered The Essential Downstream Pathways

In order to reveal the molecular mechanisms affected by *Yap1* down-regulation, proteomics was performed for Yap-MO and control-MO administered groups. Comparison of gene expression levels in blastema at 16dpa resulted in 903 identified and 285 differentially expressed (DE) proteins (Table S2). Proteomics results were validated by RT-qPCR (Fig. 3A). The observed correlation between proteomics and RT-qPCR results prompted us to continue with downstream analyses. Among these proteins, 472 (208 significant) proteins were upregulated in the Yap-MO group, whereas 431 (77 significant) genes were downregulated. Alpha-2-HS-glycoprotein (AHSG), regulator of G-protein signaling 18 (RGS18), and complement component C3 (*C3*) genes were the top significantly upregulated genes, while peptidyl-prolyl cis-trans isomerase (FKBP2), Glutathione S-transferase M1 (GSTM1), and ATPase H+ Transporting V1 Subunit G1 (ATP6V1G1) genes were detected as top downregulated genes. DE proteins were used to enrich the GO terms, KEGG pathways, and GSEA terms.

**Fig. 3.**
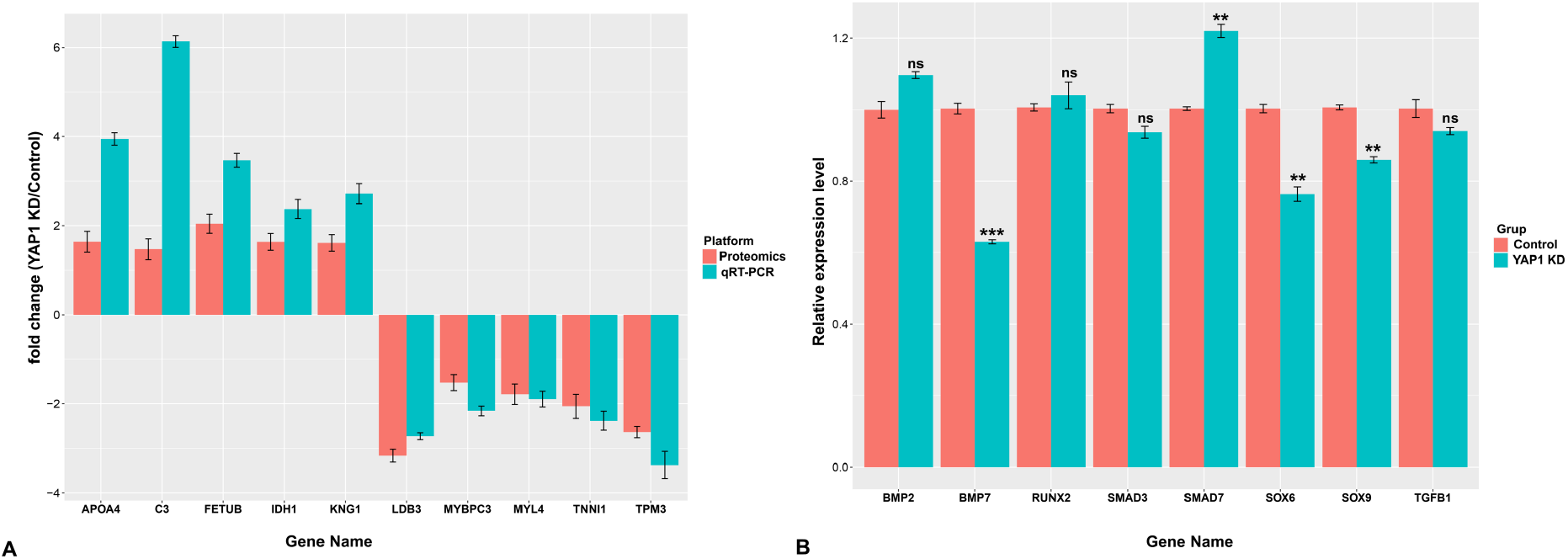
Expression levels of selected genes. Among the differentially expressed genes, 10 of them were selected for validation of proteomics. Red and blue bars show the fold change based on proteomics and qRT-PCR results, respectively (A). Expression level of osteogenesis and chondrogenesis related genes was compared between control and Yap1_KD group (B). *p-value < 0.05, **p-value < 0.01, ***p-value < 0.001, ns; non-significant. *Apoa4:* Apolipoprotein A4, *Bmp2:* Bone morphogenetic protein 2, *Bmp4*:Bbone morphogenetic protein 4, C3: Complement component 3, *Idh1:* Isocitrate Dehydrogenase (NADP(+)) 1, *Kng1:* Kininogen-1, *Ldb3:* LIM domain binding 3, *Mybpc3:* Myosin-binding protein C 3, *Myl4:* Myosin Light Chain 4, *Prdx*2:Peroxiredoxin 2, *Runx2:* Runt-related transcription factor 2, *Smad3:* SMAD family member 3, *Smad7:* SMAD family member 3, *Sox6:* SRY-Box transcription factor 6, *Sox9*: SRY-Box transcription factor 9, *Tgfb1:* Transforming growth factor beta 1, *Tnni1:* troponin I type 1, *Tpm2:* Tropomyosin 2

Top BP terms enriched by upregulated DE proteins such as ‘regulation of endopeptidase activity’, ‘regulation of peptidase activity’, and ‘negative regulation of hydrolase activity’ were related to enzymatic activities (Fig. 4A, Table S1). Upregulated DE proteins enriched ‘blood microparticle’, ‘secretory granule lumen’, and ‘cytoplasmic vesicle lumen’ CC terms, while ‘enzyme inhibitor activity’, ‘endopeptidase inhibitor activity’, and ‘peptidase inhibitor activity’ were the enriched top MF terms (Fig. 4A). On the contrary, ‘muscle contraction’, ‘muscle system process’, and ‘actin filament-based movement’ BP terms were detected for the downregulated DP proteins (Fig. 4B, Table S2). ‘contractile fiber’, ‘myofibril’, and ‘sarcomere’ were the enriched top CC terms, whereas we identified ‘actin binding’, ‘actin filament binding’, and ‘extracellular matrix structural constituent’ MF terms as the utmost significant terms (Fig. 4B). Enriched immune system-related BPs including ‘regulation of humoral immune response’, ‘neutrophil activation involved in immune response’, ‘humoral immune response’, ‘acute inflammatory response’, and ‘complement activation’ were also noticeable (Table S1, Fig. 3C). Moreover, it is noteworthy to observe that up-and down-regulated DE proteins enriched ‘muscle contraction’, ‘actin-myosin filament sliding’, ‘muscle filament sliding’, ‘actin mediated cell contraction’, and ‘actin filament based movement’ BPs (Fig. 4D, Table).

**Fig. 4.**
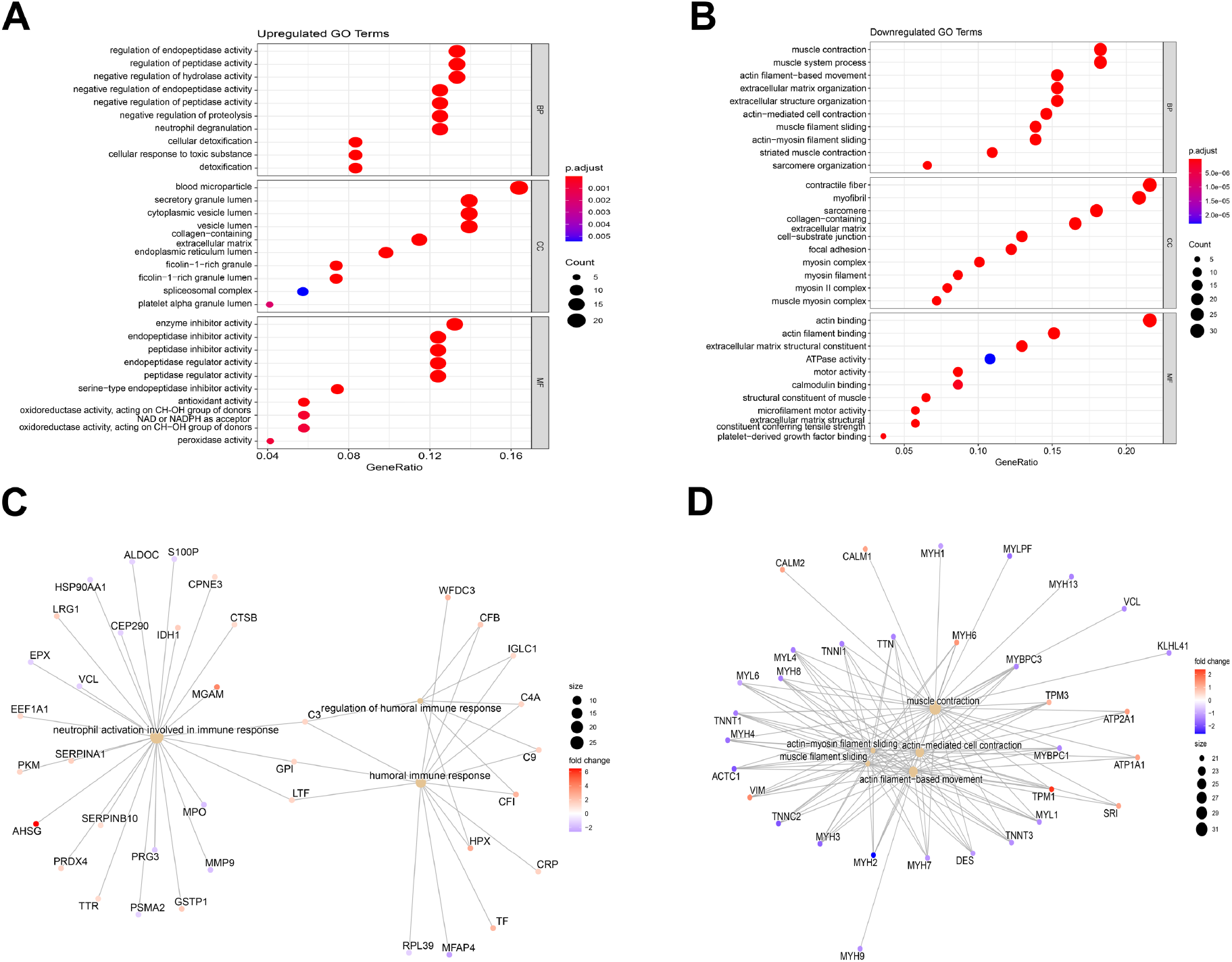
Gene ontology analysis of differentially expressed proteins. (A) top 10 GO terms enriched by 208 upregulated proteins in *Yap* depleted samples compared to the control group. (B) top 10 GO terms enriched by 77 down regulated proteins in *Yap* depleted samples compared to the control group. (C) Gene-concept network of the top 3 immune-system related biological processes enriched by significantly upregulated proteins in *Yap* depleted groups. (D) Gene-concept network of the top 5 muscle system related biological processes enriched by significantly downregulated proteins in *Yap* depleted groups.

As a subsequent analysis, KEGG pathways were enriched by the Yap-MO upregulated and downregulated DE proteins (Fig. 5A-B, Table S4-5). ‘Complement and coagulation system’, ‘glutathione metabolism’, and ’carbon metabolism’ were detected as top KEGG pathways enriched by upregulated DE proteins. On the other hand, downregulated DE proteins enriched the ‘focal adhesion’, ‘hypertrophic cardiomyopathy’, and ‘dilated cardiomyopathy’ KEGG pathways.

**Fig. 5.**
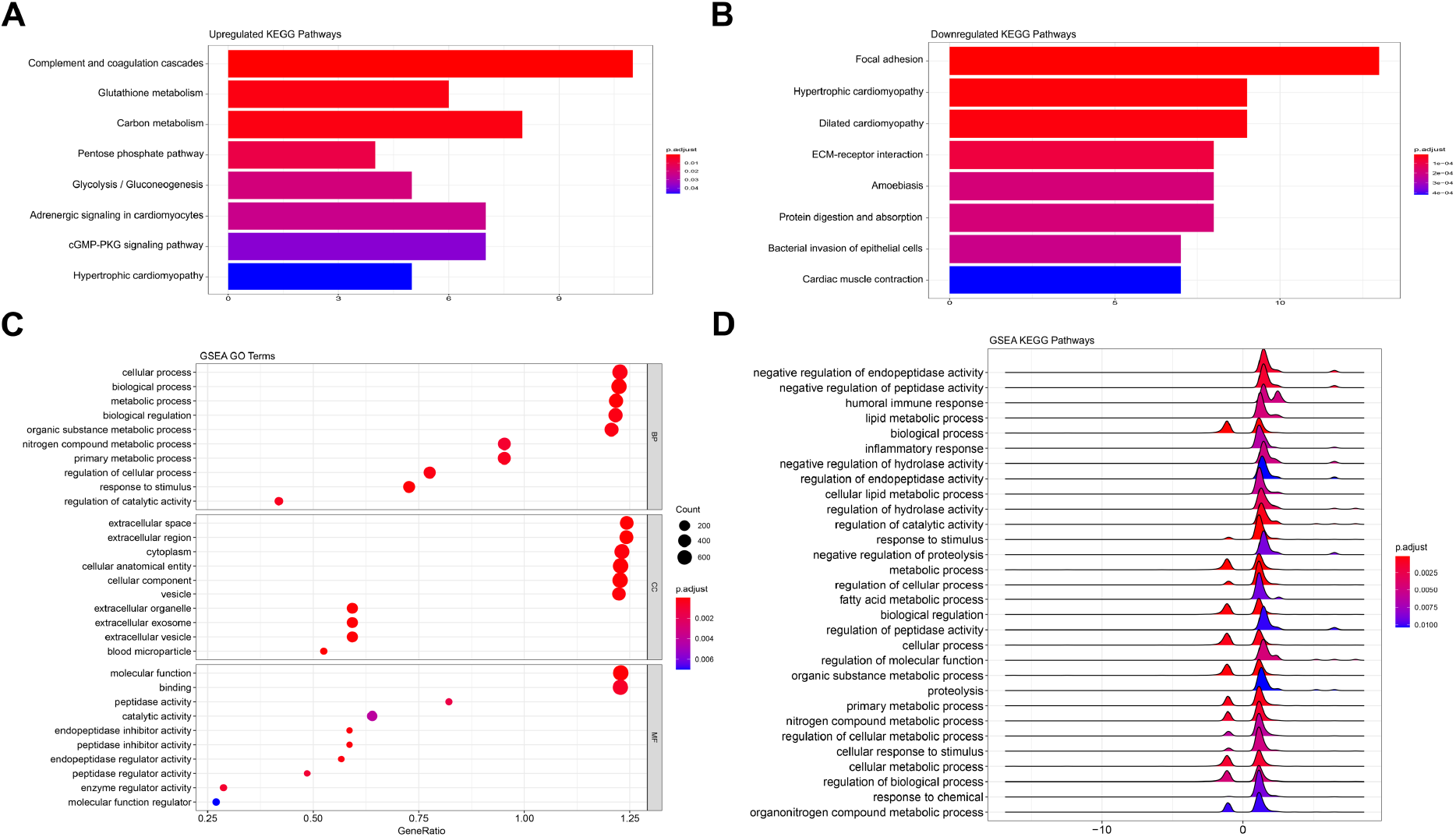
KEGG and gene ontology analyses of identified proteins. (A) Enriched KEGG pathways by upregulated proteins in *YAP* knocked-down samples compared to control group. (B) Enriched KEGG pathways by downregulated proteins in *YAP* depleted samples compared to control group. (C) top 10 GO terms enriched by identified proteins using the gene set enrichment analysis. (D) KEGG pathways enriched by the protein list using the gene set enrichment analysis.

Next, GO terms, and KEGG pathways were enriched using the GSEA method (Fig. 5C-D, Table S6). Top BPs were unveiled as ‘Cellular process’, ‘biological process’, and ‘metabolic process’ terms. ‘extracellular space’, ‘extracellular region’, and ‘cytoplasm’ were the top enriched CC terms, while the detected top MF terms were ‘molecular function’, ‘binding’, and ‘peptidase activity’. KEGG pathways were shown on a ridgeplot and ‘negative regulation of endopeptidase activity’, ‘negative regulation of peptidase activity’, ‘biological process’, ‘humoral response’, ‘inflammatory response’, and ‘lipid metabolic process’ were the top enriched KEGG pathways (Fig. 5D). The enrichment of peptidase activity inhibition, and immune system-related pathways in the Yap-MO gene set was remarkable. GSEA based enriched GO terms, and KEGG pathways were in line with DE proteins enriched results.

Lastly, the expression levels of the key genes functioning in osteogenesis or chondrogenesis, which did not appear in our proteome data, were evaluated by qRT-PCR (Fig. 3B). Among the analyzed genes, *Sox9, Sox6*, and *Bmp7* were found as significantly downregulated, and *Smad7* was found to be significantly upregulated in the *Yap1* depleted group. No significant expression level differences were detected for *Tgfb-1, Bmp2*, and *Runx2* genes between *Yap1*_KD and control groups.

## 4. DISCUSSION

Axolotl is an emerged animal model to dissect the molecular mechanisms underlying regeneration (McCusker and Gardiner, 2011). It holds great promise to be utilized in regenerative medicine studies due to its exceptional regenerative potential and advantageous experimental features (McCusker et al., 2015). In this study, we interrogated the putative regulatory roles of *Yap1* in axolotl limb regeneration. YAP, as one of the effector proteins of the hippo pathway, plays a key role in organ growth, regeneration of tissues, and many other cellular regulations such as proliferation and differentiation (Kovar et al., 2020). The amount of YAP and its phosphorylation status are important to determine the fate of cells (Moya and Halder, 2018).

Therefore, first, *Yap1* mRNA level was compared between metamorphic and neotenic animals since the previous studies reported diminished regeneration ability after metamorphosis (Demircan et al., 2018; Monaghan et al., 2014). Significantly higher *Yap1* mRNA and protein levels at 10dpa in blastema tissue of neotenic axolotls compared to metamorphic animals highlighted that increased *Yap1* expression might facilitate the successful regeneration process. Our finding supported the earlier studies conducted with invertebrate and vertebrate models, which disclosed the necessity of increased *Yap1* levels during regeneration (Hayashi et al., 2014b). Observed expression level differences prompted us to inhibit the *Yap1* activity in the blastema stage. Administration of Yap-MO decreased the size of the blastema area, and *Yap1* inhibition led to defects in bone regeneration. It was highly remarkable to obtain abnormal bone structure and missing bones when *Yap1* is depleted during limb regeneration. Previously published reports strongly support this study’s main finding, which links bone formation deficiencies to *Yap* downregulation. As documented earlier, the *Yap1* level and its interactions are vital for differentiating human mesenchymal stem cells to chondrocytes and osteoblasts (Li et al., 2021). One of the pioneering studies aiming at deciphering *Yap* activity found that a constitutively active *Yap* mutant overexpression promoted osteogenic differentiation, and osteogenesis is inhibited upon depletion of *Yap* (Dupont et al., 2011). Subsequent studies further confirmed the importance of *Yağ* in bone formation and repair. It has been demonstrated that YAP1 is required for osteoblast progenitor cell proliferation and OB differentiation to promote osteogenesis and maintain bone hemostasis through the Wnt/β-catenin signaling pathway (Pan et al., 2018). In that study, *Yap1* is conditionally inhibited in young adult mice, and this genetic manipulation resulted in bone loss (Pan et al., 2018). In another recent study, increased osteoclastic activity and, therefore, a diminished osteoblastic activity was described due to *Yap1* knockout in mice (Kegelman et al., 2018). Decreased osteogenic and collagen-related gene expression as a consequence of *Yap/Taz* gene deletion and inhibition of *Yap/Taz* with its transcriptional coeffector TEAD (Kegelman et al., 2018) provided another layer of evidence on YAP1 regulation in bone formation. A similar conclusion was achieved by Lorthongpanich and his colleagues in which gain- and loss-of-function experiments validated YAP’s essential roles in osteogenic differentiation (Lorthongpanich et al., 2019). Altogether, the observed osteogenic defects upon *Yap1* depletion during axolotl limb regeneration ties well with the established roles of YAP1 in the proliferation of OB progenitors and OB differentiation.

Besides the functions of YAP/TAZ proteins in bone development in mammals (Kovar et al., 2020), their essential role in Xenopus regeneration has been explored as well (Hayashi et al., 2014a; Hayashi et al., 2014b). Dominant-negative *yap1 (dnyap1*) activity during Xenopus limb regeneration affected crucial biological processes such as cell death and cell proliferation, and as a consequence of wild-type *yap1* inhibition, a decrease in bone length accompanied by digit formation reduction were observed (Hayashi et al., 2014b). In another study, Hayashi et al., followed a similar experimental design to interrogate the roles of *yap1* and *tead4* in tadpole tail regeneration (Hayashi et al., 2014a). The size of the regenerated tail in transgenic frogs expressing *dnyap1* was shorter than the control group, and a small proportion of dnyap1 animals could regenerate their missing tails (Hayashi et al., 2014a). Observation of incomplete regeneration or severely defective regeneration due to increased apoptosis and decreased proliferation rate in some *dnyap1* animals (Hayashi et al., 2014a) highlighted the necessity of *yap1* in appendage regeneration and organ growth. Our results are well-aligned with findings in Xenopus, underlining the conserved activity of *yap1* among animals in regeneration.

Furthermore, our data on the necessity of YAP1 for a successful regeneration confirmed a previous study conducted in axolotl. It has been reported that *Yap1* expression and activation by Hydrogen peroxide is necessary for successful tail regeneration (Carbonell et al., 2021). Reactive oxygen species generated during regeneration regulate immune cells’ recruitment to the wound zone and activation of Akt and YAP signalling pathways to positively regulate the cell cycle and survival (Carbonell et al., 2021). Administration of *Yap1* inhibitor during tail regeneration resulted in defects in appendage renewal, implying the indispensable role of *Yap1* in epimorphic regeneration.

We then sought the downstream pathways affected due to the *Yap1* knock-down by employing the proteomics method to provide a mechanistic explanation. The altered expression level of muscle-activity-related genes at the blastema stage of axolotl limb regeneration through dedifferentiation of specialized cells into the stem and progenitor cells has been reported earlier (Rao et al., 2009; Sibai et al., 2020).Interestingly, in our dataset, several genes with muscle activity such as *Tpm1, Tpm3, Myh6*, and *Myh7b* have been found upregulated, whereas the genes such as *Tnnc1, Tpm2, Myo2*, and *Myh1* were downregulated following the *Yap1* depletion, highlighting an ambiguous effect of *Yap1* loss on muscle activity at the early stage of limb regeneration. Enrichment of the BPs related to negative regulation of peptidase activity by the upregulated genes in Yap1_KD samples was noteworthy. Our results are in accordance with earlier investigations which showed that peptidase activity is required for an appropriate regeneration process in both invertebrates and vertebrates (Dolmatov et al., 2019; Dong et al., 2021; Enos et al., 2019; Pasten et al., 2012; Rinkevich et al., 2007).

A considerable decrease in the regenerated intestine of the sea cucumber was identified following the inactivation of peptidases by administration of inhibitors (Pasten et al., 2012). MG132 (a reversible chymotrypsin and peptidylglutamyl peptidase hydrolase inhibitor), E64d (a cysteine proteases inhibitor), and TPCK (a serine chymotrypsin inhibitor) interfered with successful regeneration process due to the impact of peptidase inhibition on cell proliferation, extracellular matrix remodeling, and apoptosis (Pasten et al., 2012). Furthermore, Urochordate ascidians’ whole body regeneration is impaired with serine protease inhibition due to the disruption of the vascular environment (Rinkevich et al., 2007). Decreased activity of proteases accounted for the formation of a dense scaffold by deposition of cellular and matrix materials to the wound site, and developed disorganized structures failed to regenerate (Rinkevich et al., 2007). In holothurians, inhibition of proteinase activity at the early time point of regeneration abolished the regeneration process, whereas inhibition at late time points delayed the ambulacral structures regeneration (Dolmatov et al., 2019). On the other hand, axolotl Cathepsin K (a cysteine protease) and ependymal MMPs take a role in ECM degradation during spinal cord regeneration (Enos et al., 2019). Therefore, we can conclude that, the link between decreased regeneration fidelity upon *Yap1* knock-down and detected over-represented BPs related to negative regulation of peptidase, endopeptidase, hydrolyse and proteolysis activity is broadly consistent with previous reports.

Moreover, upregulated genes in *Yap1* knocked-down samples enriched immune system BPs such as ‘neutrophil degranulation’, ‘neutrophil activation involved in immune response’, and ‘acute inflammatory response’ were worth mentioning. The immune system is vital for a proper regeneration process (Julier et al., 2017). However, the prolonged and over-activated immune system response may cause the scarring and prevent the regeneration program (Aurora and Olson, 2014). Single-cell RNA sequencing study in axolotl unveiled that neutrophils infiltrate to the wound bed within 24h of amputation, and their abundance decreases substantially at 6dpa (Rodgers et al., 2020). Hence, increased neutrophil activity in the blastema of the *Yap1* knocked-down samples might be one of the reasons for the delayed and incomplete regeneration. Furthermore, a proteomics study aimed at comparison of gene expression profile during limb regeneration of neotenic and metamorphic axolotl described a strong immune response in blastema tissue of regeneration deficient metamorphic animals (Sibai et al., 2020). In another study, transcriptome analysis during the spinal cord regeneration in metamorphic axolotl was performed and a prolonged immune system activity at the blastema stage was detected (Demircan et al., 2020). It was suggested that the observed delay in metamorphic axolotl spinal cord regeneration might be linked to extended immune response. In zebrafish heart regeneration, it has been exhibited that enhancing activity of *yap1-ctgfa* signaling during cardiac regeneration is also due to negative regulation of macrophage migration and infiltration into the injury site (Flinn et al., 2019; Mukherjee et al., 2021). Likewise, defective heart regeneration in *yap1* knock-out fish was linked to an increased level of scarring because of the high rate of macrophage infiltration to the wound site (Mukherjee et al., 2021).

Additionally, the roles of several DE proteins in bone formation have been identified previously. Cathepsin K (CTSK), a matrix-protease, is required for osteocyte remodeling and bone homeostasis, and YAP positively regulates its expression (Kegelman et al., 2020). *Yap1* knock-down significantly and considerably decreased its level, which is consistent with its regulation by YAP1. Mice with conditionally inactivated *Yap/Taz* exhibited a reduced expression level of CTSK, and the percentage of CTSK positive osteocytes decreased significantly (Kegelman et al., 2020). The bone healing activity of FXIII has been revealed (Reviewed in (Kleber et al., 2022)), and the diminished level of FXIII protein after *Yap1* inhibition might be another contributor to the observed bone defects. Similarly, the apolipoprotein-E level was dropped in *Yap1* depleted animals and considering its role in suppressing osteoclastogenesis (Kim et al., 2013; Niemeier et al., 2012), it is plausible to suggest that a low level of APO-E results in increased osteoclast differentiation and this may further explain the deficient regeneration process. Moreover, a component of vacuolar ATPase (ATP6V1H) was another downregulated DE protein in our list, and based on the literature, its deficiency caused increased bone resorption and decreased osteoblast activity (Duan et al., 2016; Zhang et al., 2017).

Also, dysregulation in ECM components, ECM remodeling enzymes, and intracellular elements of cell-cell, cell-ECM interaction complexes by *Yap1* inhibition in axolotl limb regeneration is worth mentioning. As shown previously, ECM stiffness is directly related to lineage specification and a key regulator of YAP/TAZ subcellular localization and activity (Dupont et al., 2011; Engler et al., 2006; Han et al., 2019). Many proteins and glycoproteins with structural and signaling roles in ECM and intracellular network, such as several collagen types, tenascin, transforming growth factor beta-induced (TGFBI), decorin and vinculin were downregulated in *Yap1* depleted group compared to the control. An extracellular matrix glycoprotein, Tenascin C (TNC), plays a fundamental role in new bone formation through the activation of the TGF-β signaling pathway, and its expression is elevated during osteogenesis (Li et al., 2016; Sato et al., 2016). In bone development, the roles of TGFBI, an ECM protein that is strongly induced by TGF-β (Thapa et al., 2007), have been revealed in a previous study that reported the reduced bone mass as a result of *Tgfbi* gene disruption (Yu et al., 2012b). As a small leucine-rich proteoglycan (SLRP), decorin is an essential proteoglycan in the bone with cell proliferation, osteogenesis, mineral deposition, and bone remodeling roles during bone formation (Kirby and Young, 2018). Stiffness of ECM affects the level and localization of vinculin, a major regulator of the focal adhesion pathway, and decreased level of vinculin or its localization in the nucleus reduces osteoblast differentiation (Zhou et al., 2019).

Last but not least, the expression level of the primary regulators of cartilage and bone formation was investigated. A significant decrease in *Sox9, Sox6*, and *Bmp7* through *Yap1* inhibition is in accordance with previous studies describing a direct upregulation of these genes by the YAP1-TEAD binding to the target promoters (Park et al., 2015; Song et al., 2014). Genome-wide cooperation of SOX9 with SOX6, mainly through binding to enhancers to regulate the chondrocyte differentiation program, has been demonstrated before (Liu and Lefebvre, 2015). Although BMPs have a strong osteogenic capacity, due to the functional redundancy, the effect of BMP7 depletion on cartilage development was found to be minor (Shu et al., 2011). The increased level of inhibitory *Smad7* in the *Yap1* depleted group is noteworthy. It has been disclosed that SMAD7 negatively regulates chondrocyte proliferation and chondrocyte maturation by interfering with BMP and TGFβ signaling pathways (Iwai et al., 2008; Moustakas and Heldin, 2009). Taken together, our findings pinpoint that the expression level of several genes with chondrogenic and osteogenic activities are altered by decreased *Yap1* expression; however, deciphering the exact mechanism requires further investigation.

It is tempting to speculate that one of these genes or a combinatory effect led to bone formation defects; however, a detailed characterization of these candidate genes’ roles in limb regeneration is necessary to draw a precise conclusion. One of the deficiencies of the presented study is the lack of *Yap1* over-expression data in limb regeneration which would have contributed more to elucidating its roles during functional restoration. In addition, an earlier and extended knock-down of *Yap1* during regeneration would be vital for a full dissection of its regulatory functions during limb renewal.

Overall, the present study provides new evidence on the defects in regeneration due to Yap1 depletion. Our data indicated that bone formation during axolotl limb regeneration in *Yap1* knocked-down animals was probably hampered by increased immune system activity, negative regulation on peptidases, changed ECM composition, and the altered expression level of the proteins required for a successful osteogenesis.

## 5. CONCLUSION

In this study, the role of YAP1 protein in axolotl limb regeneration was examined. We demonstrated that the *Yap1* expression level is higher in neotenic axolotls than in the metamorphic animals at the early stages of limb regeneration. Furthermore, the indispensable role of YAP1 in limb regeneration was evident by bone formation defects as a consequence of *Yap1* depletion. BPs affected by *Yap1* inhibition were interrogated employing the proteomics method. Enrichment of immune system-related, and enzymatic activity inhibitory pathways by differentially upregulated proteins in Yap-MO injected samples might explain the observed defects in axolotl limb regeneration. Follow-up research to modulate the expression level or activity of the candidate proteins described in this study during axolotl limb regeneration may bridge the gap between the observed infidel bone regeneration and altered levels of the proteins.

## Supporting information

Primer sequences to amplify Yap1 and Elf1&#945; (housekeeping gene used for normalization) were provided in Table S1.

blastema at 16dpa resulted in 903 identified 359 and differentially expressed (DE) proteins (Table S2)

## AUTHOR CONTRIBUTIONS

S.B.: Provision of study material, Collection and assembly of data, Data analysis and interpretation, Manuscript writing, Final approval of manuscript; G.Ö.: Conception and design, Manuscript writing, Final approval of manuscript; N.E.: Conception and design, Manuscript writing, Final approval of manuscript; T.D.: Conception and design, Provision of study material, Collection and assembly of data, Data analysis and interpretation, Manuscript writing, Final approval of manuscript.

## ACKNOWLEDGEMENTS

We thank Ecem Yelkenci, Elif Özbay, Pelin Tuğlu, and Sultan Gül for their help in proteomics experiments and animal care.

